# Comprehensive analysis of full-length transcripts reveal aberrations of splicing variants in liver cancer

**DOI:** 10.1101/2021.06.28.450266

**Authors:** Hiroki Kiyose, Hidewaki Nakagawa, Atsushi Ono, Hiroshi Aikata, Masaki Ueno, Shinya Hayami, Hiroki Yamaue, Kazuaki Chayama, Mihoko Shimada, Jing Hao Wong, Akihiro Fujimoto

## Abstract

Genes generate various transcripts by alternative splicing, and these transcripts can have diverse functions. However, in most transcriptome studies, short-reads sequencing technologies (next-generation sequencers) have been used and full-length transcripts have not been observed directly. Although long-reads sequencing technologies would enable us to sequence full-length transcripts, analysis of the data is a difficult task. In the present study, we developed an analysis pipeline named SPLICE to analyze full-length cDNA sequences. Using this method, we analyzed cDNA sequences from 42 pairs of hepatocellular carcinoma (HCC) and matched non-cancerous liver with Oxford Nanopore technology. Our analysis detected 46,663 transcripts from the protein-coding genes in the HCCs and the matched non-cancerous livers, of which 5,366 (11.5 %) were novel. Comparison of expression levels identified 9,933 differentially expressed transcripts (DETs) in 4,744 genes. Importantly, 746 genes with DET were not found by the gene-level analysis. We also identified novel exons derived from transposable elements (TEs). In the analysis of transcripts from hepatitis B virus (HBV), HBx-human TE fusions were found to be overexpressed in the HCCs. Furthermore, fusion gene detection showed novel recurrent fusion events. These results suggest that long-reads sequencing technologies allow us to analyze full-length transcripts, and show the importance of splicing variants in carcinogenesis.

## Introduction

Hepatocellular carcinoma (HCC) is the third-leading cause of death worldwide and the seventh most common form of cancer (Bray et al. 2018). Common etiological factors in liver carcinogenesis include infection by hepatitis B virus (HBV), and hepatitis C virus (HCV), but other factors, such as alcohol intake, metabolic diseases, and specific carcinogen exposures also play significant roles (El–Serag and Rudolph 2007). These factors cause liver inflammation, leading to cirrhosis and resulting in malignant transformations in hepatocytes (El–Serag and Rudolph 2007). To elucidate the molecular mechanisms underlying liver carcinogenesis, genetic and transcriptional aberrations have been investigated (Ally et al. 2017; Fujimoto et al. 2016; Guichard et al. 2012; Schulze et al. 2015; Totoki et al. 2011). These previous studies identified numerous somatic mutations and differentially expressed genes, and have led to the discovery of liver cancer-associated pathways, such as apoptosis, Wnt signaling, chromatin remodeling and lengthening telomeres (Ally et al. 2017; Fujimoto et al. 2016; Guichard et al. 2012; Schulze et al. 2015; Totoki et al. 2011). In HBV-related liver cancers, HBV integrations and HBV-human fusion transcripts have been identified (Shiraishi et al. 2014). Fusion genes have been investigated, but recurrent fusions were rare (Vellichirammal et al. 2020). Although these studies have expanded our knowledge on HCC, our understanding of liver carcinogenesis is still far from complete (Alqahtani et al. 2019; Liu et al. 2015). To obtain deeper insight into the mechanism responsible for HCC, applications of novel technologies and analysis of new aspects of genetic and transcriptional aberration would be needed.

We aimed to analyze full-length transcripts for HCC in the present study. The majority of previous transcriptome studies have used microarrays or short-read sequencing technologies, and therefore lacked information on full-length transcripts. Analysis of full-length transcripts should enable us to detect splicing variants expressed from each gene. Protein sequences are specified by mRNA sequences of transcripts, thus direct observation of transcripts should provide us with essential information about carcinogenesis. Indeed, several recent studies have suggested that transcript-specific functions contribute to carcinogenesis such in breast and ovarian cancers (David and Manley 2010; Venables 2004; Venables et al. 2009). In HCC, a recent study analyzed short-read RNA-seq data and detected oncogenic splicing changes in *AFMID* gene (Lin et al. 2018). These splicing changes were estimated to occur at the early stage of HCC carcinogenesis, and were associated with the patients’ survival (Lin et al. 2018). These previous studies strongly suggest that splicing variants have important roles, most of which would have been missed, and therefore, analysis of full-length transcripts should be significant for cancer studies.

One promising approach to observe transcripts is RNA-seq using long-read sequencers (Sakamoto et al. 2020), however, their application in the cancer research field is still limited (Bueno et al. 2016; Huang et al. 2021; Lian et al. 2019; Oka et al. 2021; Tang et al. 2020), likely due to high error rates of long-read sequencing (Rang et al. 2018) and complexity or heterogeneity of cancer genome. To overcome this problem and identify cancer-related transcriptional changes, we developed a method for analyzing RNA-seq data obtained by a long-read sequencer MinION (Oxford Nanopore). In this study, we sequenced 42 pairs of cDNA from HCCs and matched non-cancerous livers, which have been previously used for whole-genome sequence (WGS) and RNA-seq by short-reads (Fujimoto et al. 2016; Fujimoto et al. 2012; ICGC/TCGA Pan-Cancer Analysis of Whole Genomes Consortium 2020). We developed an analysis method named SPLICE and analyzed the long-read data. Among reads mapped to coding genes, 61.7 % contained entire cording sequences, and a total of 46,663 transcripts were detected in the protein-coding genes in the HCCs and matched livers, of which 5,366 (11.5 %) were novel. Comparison of expression levels found 9,933 differentially expressed transcripts (DETs) in 4,744 genes. Interestingly, 746 genes with DET were not detected by the gene-level analysis. Exonization of transposable elements (TEs) was also identified. In the analysis of HBV transcripts, fusion transcripts of HBV HBx gene and human TEs were overexpressed in HBV-related HCCs. These results suggest that long-reads sequencing technologies enable us to gain a fuller understanding of cancer transcripts, and the SPLICE method contributes to the analysis of transcriptome sequences by the long-reads sequencing technology.

## Results

### cDNA sequencing with Oxford Nanopore sequencer

We sequenced cDNA from MCF-7 cell, and previously published 42 cancer and their matched non-cancerous liver pairs (**Supplemental Table S1-4**; Fujimoto et al. 2016). cDNA was generated by reverse-transcription with SMARTer PCR cDNA synthesis kit (TakaraBio), and sequencing libraries were constructed using the SQK-LSK109 library construction kit (Oxford Nanopore) according to manufacturer’s instruction. We performed one run for each sample with flowcell R9.4 (Oxford Nanopore). The data yields ranged from 4.6 Gbp to 20.6 G bp (average 12.7 Gbp). The number of reads were 3,061,266 to 14,873,634 (average 9.1 k reads) (**Supplemental Table S4**).

### Construction of analysis pipeline and evaluation of the SPLICE method

To analyze long-read data, we constructed an analysis pipeline named SPLICE (**see Methods**; **Fig. 1A**). Long-read sequencers have the capability to sequence entire cDNA end-to-end (Bolisetty et al. 2015), allowing us to capture full-length transcripts without assembly (Trapnell et al. 2010). However, long-read sequencers have high error rates (Rang et al. 2018) and would cause errors in identifying splicing variants and estimating their expression levels. To overcome these obstacles, we analyzed error patterns and developed an analysis pipeline.

**Fig. 1:**
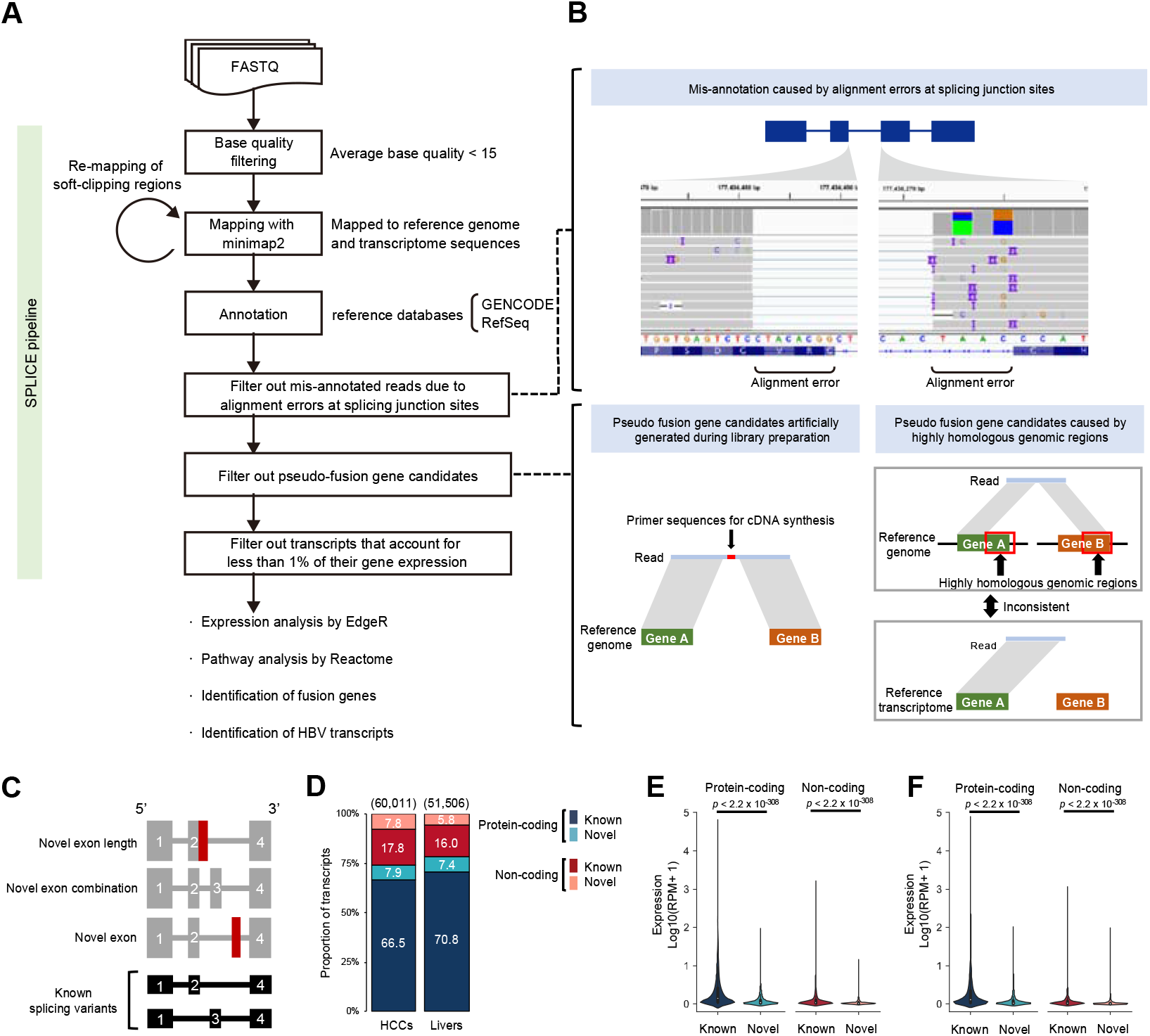
Overview of the SPLICE pipeline and HCC sequencing library. (**A**) Workflow of the SPLICE pipeline and data analysis of this study. Reads with an average base quality score < 15 were removed from the input FASTQ files. The read sequences were mapped to the reference genome (hg38) and the reference transcriptome sequences (GENCODE (version 28) and RefSeq (release 88)), and remapped soft-clipped regions (≧ 60 bp). Candidates corresponding to the error patterns or low-frequency transcripts were removed. The final data were used for subsequent data analyses. (**B**) Error patterns in cDNA-seq using MinION. Alignment errors caused by sequencing errors, artificial chimeras, and mapping errors of highly homologous genes can result in errors. (**C**) Classification of patterns of novel transcripts. “Novel exon length” contains known exons with different lengths from the reference database. “Novel exon combination” has a different exon combination from the reference database. “Novel exon” contains exons expressed from regions that are not overlapped with known exons. (**D**) Composition of the transcripts. The stacked bar chart shows the distribution (%) of detected transcripts by transcript type. The total number of transcripts is shown above the bars. (**E,F**) Comparison of expression levels of known and novel transcripts. Transcript abundance was measured in reads per million reads (RPM) and log10 converted values for RPM+1 were shown in the violin plot. *P*-values were calculated by Wilcoxon rank-sum test. (**E**) Comparison in HCC samples. (**F**) Comparison in non-cancerous liver samples.

Reads were mapped to the reference genome sequence (hg38) and reference transcriptome sequence (GENCODE (version 28) and RefSeq (release 88)) with minimap2 software (Li 2018). We found that false-positive splicing aberrations were caused by alignment errors at splicing junction sites, mapping errors due to highly homologous genomic regions, and artificial chimeric reads (**Fig. 1B**). The following describes how the SPLICE pipeline removed possible error patterns from the mapping results; (1) Reads were mapped to the reference genome and the reference transcriptome sequences, and reads were removed if the number of matched bases in a read to the reference genome sequence was smaller than that to the reference transcriptome sequence (removal of mapping errors). (2) In the identification of novel transcripts (novel splicing variants), we evaluated the error rate within the ± 5 bp regions from splicing junction sites and removed candidates if error rates ≧ 20 % (removal of alignment errors around splicing sites). (3) Fusion gene candidates can be generated by mis-ligation of two transcripts during library preparation, we therefore removed fusion gene candidates that contained primer sequences for cDNA synthesis between two genes (removal of artificial chimeras). (4) Mapping errors due to highly homologous genomic regions also can cause fusion gene candidates, thus, we compared the results of mapping to the reference genome and the reference transcriptome sequences, and removed candidates if both were inconsistent (removal of mapping errors). (5) We removed transcripts if their proportion of expression level of a transcript was less than 1 % of the total amount of expression of a gene (removal of possible artificial chimeras). Using these filters, we estimated the expression level of transcripts and identified novel transcript candidates from long-reads, and classified them into transcript with “novel exon length”, “novel exon combination”, and “novel exon” (**Fig. 1C**).

To evaluate this method, we sequenced cDNA obtained from MCF-7 cell line, and analyzed them with the SPLICE pipeline. As a result of the analysis, 13,424,802 reads (99.5 %) were mapped (**Supplemental Table S2**). We calculated the number of reads mapped to each transcript per 1 million reads to estimate the expression levels. Then the numbers of reads of each transcript were summed by each gene and compared to the expression level by short-read sequencing technology (ENCODE Project Consortium 2012). The gene expression levels between long-reads and short-reads showed a strong correlation (*p* < 2.2 × 10^-308^, *r* = 0.80) (**Supplemental Fig. S1A**). We then evaluated the sensitivity in detecting fusion genes. Previous studies reported 5 highly-expressed fusion genes in the MCF-7 cell (Edgren et al. 2011; Hampton et al. 2008), and our analysis identified reads for all of them (**Supplemental Table S3**), although two of them were supported by one or two reads. Based on these results, we considered that our analysis pipeline has sufficient accuracy for analyzing transcript aberrations in cancers.

### Analysis of HCC and non-cancerous liver samples

We then analyzed sequence data from the HCCs and the matched non-cancerous livers. After filtering low-quality reads (average base quality score < 15), we obtained an average of 9,043,803 reads (12.9 Gbp on average) in the HCCs and 9,013,602 reads (12.4 Gbp on average) in the livers (**Supplemental Table S4**). The average mapping rates were 98.9 % for the HCCs and 97.3 % for the livers (**Supplemental Table S4)**. Using the SPLICE pipeline, we identified transcripts supported by ≧ 3 reads and obtained a total of 60,011 non-redundant transcripts in the HCCs and 51,506 in the livers (**Fig. 1D**). Of these, 66.5 % and 70.8 % were known protein coding transcripts, 7.9 % and 7.4 % were novel protein coding transcripts, 17.8 % and 16.0 % were known non-coding transcripts, and 7.8 % and 5.8 % were novel non-coding transcripts in the HCCs and the livers, respectively (**Fig. 1D**). We then compared the expression levels of known and novel transcripts. The average expression levels of novel transcripts were lower than those of known transcripts for both protein-coding and non-coding transcripts in the HCCs and the livers (*p* < 2.2 × 10^-308^ for protein-coding, *p* < 2.2 × 10^-308^ for non-coding) (**Fig. 1E,F**).

Among reads mapped to known protein coding transcripts, 61.7 % had full-length coding sequences (CDSs) (**Supplemental Fig. S1B,C**). The proportion of reads mapped to the full-length CDS region was negatively correlated with the length of genes (**Supplemental Fig. S1D**). The proportions of reads containing the full-length CDS were correlated with RIN (RNA Integrity Number) values of the RNA samples (*p* = 9.8 × 10^-6^, *r* = 0.46) (**Supplemental Fig. S1E**), indicating the importance of RNA quality for identifying full-length transcripts. We then compared the gene expression levels between previous short-reads and the current long-reads data (Fujimoto et al. 2016), and found high correlations (*p* < 2.2 × 10^-308^ in all samples, average *r* = 0.79) (**Supplemental Fig. S1F**).

### Features of novel exons

We identified novel exons by comparing the exon locations in the database (GENCODE (version 28) and RefSeq (release 88)). Our analysis identified 767 and 531 novel exons in 4,769 and 3,786 transcripts of protein-coding genes in the HCCs and the matched non-cancerous livers, respectively. Most of them (92.0 %) were the first or the last exon of transcripts (**Fig. 2A**). Since transposable elements (TEs) are known to be exonized by tissue-specific or cancer-induced hypomethylation of the genome (Elbarbary et al. 2016), we calculated the proportion of TE-derived novel exons. In the analysis, novel exons that were > 70 % covered by TEs were considered as TE-derived novel exons. The proportion of TE-derived novel exons was significantly higher than that of TE-derived known exon in the first, last, and middle exons (*p* = 5.9 × 10^-153^ for the first exon, *p* = 2.2 × 10^-106^ for the last exon, *p* = 1.7 × 10^-79^ for middle exon) (**Fig. 2B**; **Supplemental Table S5**). A permutation test showed that the novel first exon was significantly enriched in L2 antisense, ERV1 antisense, and ERVL-MaLR sense (**Fig. 2C**). The novel last exon was significantly enriched in *Alu* sense, and the middle exon was significantly enriched in *Alu* antisense, ERVL-MaLR antisense, and ERV1 antisense (**Fig. 2C**). These results showed that exonized repeat families and strands differ by their position in the transcripts.

**Fig. 2:**
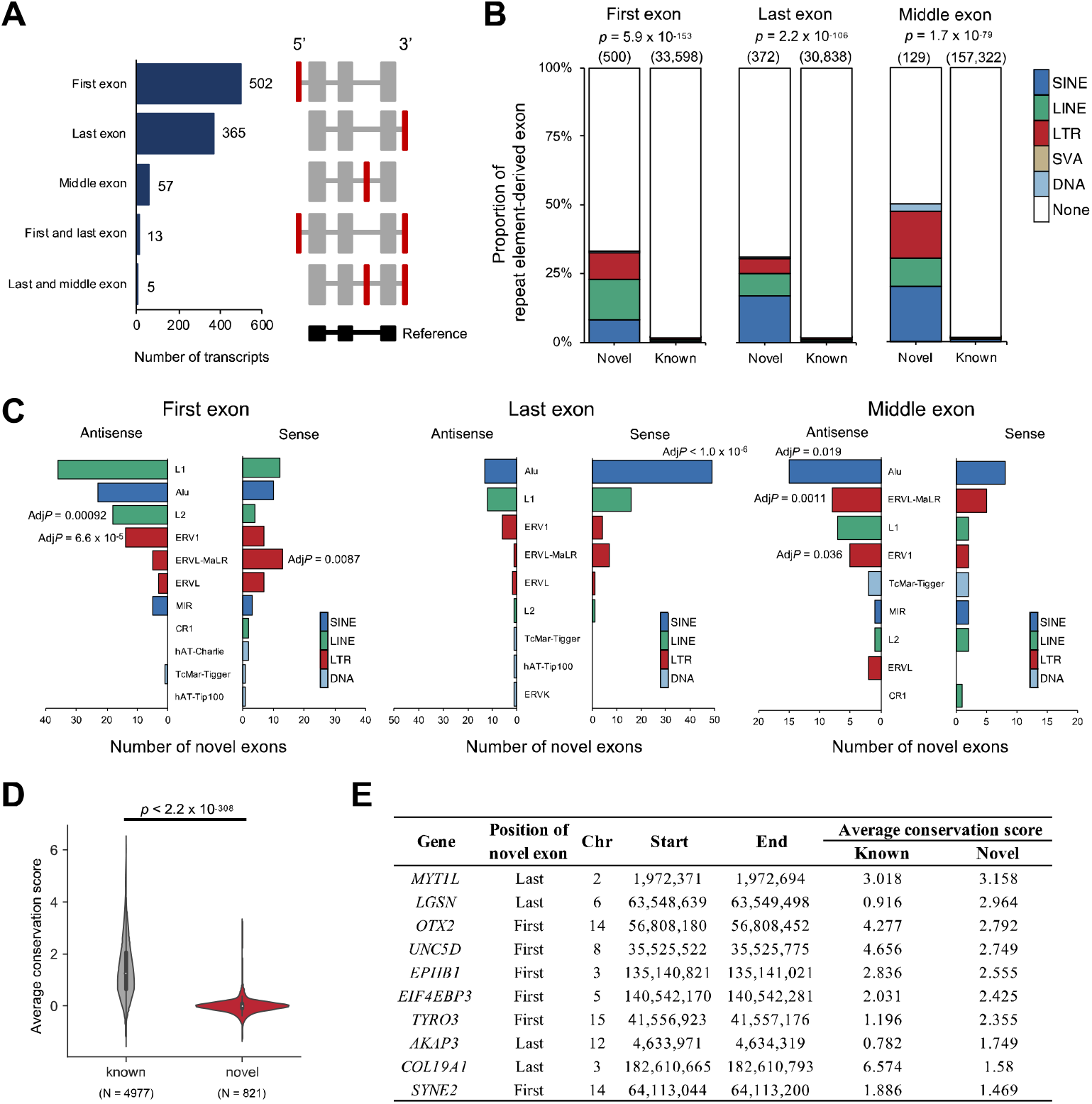
Features of novel exons. (**A**) Patterns of transcripts with novel exons. (Left panel) Bar graph showing the number of transcripts by the position of the novel exon. (Right panel) Schematic diagram of the location of the novel exons. The novel exons are shown in red. (**B**) Comparison of the proportion of TE-derived exon in novel and known exons by exon position. The stacked bar chart shows the distribution of TE-derived exons. The total number of exons is shown above the bars. Statistical significance of the total abundance of TE-derived exons between the known and novel exons was calculated by Fisher’s exact test. (**C**) The number and enrichment of TE-derived novel exons. TEs were classified by repeat family, and the sense or antisense strand for the gene strand was distinguished. The statistical significance was examined by a permutation test. *P*-values were adjusted by Bonferroni correction (Adj*P*). (**D**) Comparison of average conservation scores between novel and known exon regions. For each gene, the average conservation scores were calculated based on the phastCons scores database (100 vertebrates) (**see Data access**). *P*-values were calculated by Wilcoxon rank-sum test. (**E**) List of transcripts with novel exon with high average conservation scores. The 10 candidates were shown in order of average conservation score.

To estimate the biological importance of the novel exons, we estimated their evolutional conservation using a conservation score obtained by 100 species (**see Data access**). The average conservation score of the novel exon region was significantly lower than that of the known exon region (*p* = 2.2 < 10^-308^) (**Fig. 2D**), suggesting that most of the novel exons had less importance in the evolutional aspects. However, a part of novel exons was highly conserved (**Fig. 2E**; **Supplemental Table S6**), suggesting that they had important biological functionalities.

### Identification of significantly differentiated genes and transcripts

One of the most important features of long-read transcriptome is detecting differentially expressed transcripts. Thus, comparing the expression level between the HCCs and matched non-cancerous livers, we aimed to find differentially expressed genes (DEGs) and differentially expressed transcripts (DETs). DETs are transcripts that showed significant differentiation in expression level between the HCCs and the matched non-cancerous livers. DEGs are genes obtained by the comparison of the total amount of transcripts within a gene between the HCCs and the livers. We estimated the expression level based on the number of reads and compared them with the edgeR software (**see Methods**; Robinson et al. 2010). Our analysis identified 4,744 DEGs and 9,933 DETs after adjusting for multiple testing (FDR < 0.01) (**Fig. 3A**; **Supplemental Table S7**; Benjamini and Hochberg 1995). Reactome pathway analysis (Jassal et al. 2020) for genes with up-regulated DETs was conducted, and genes with the terms “M Phase”, “Cell cycle, Mitotic”, and “Cell cycle” were significantly enriched (**Supplemental Table S8**). The aforementioned transcript with a conserved novel exon in the *MYT1L* gene was significantly down-regulated (FDR = 0.00088) **(Fig. 2E**; **Supplemental Fig. S3A,B**; **Supplemental Table S7**). Since *MYT1L* has an antitumor effect in glioblastoma multiforme (Hu et al. 2020), this transcript would have an important role in HCCs. Among the genes with DETs, 746 genes (935 transcripts) were not detected as DEGs (DET-specific genes) (**Fig. 3B**; **Supplemental Table S7**). To know the features of the DET-specific genes, we compared the numbers of transcripts identified in this study and transcription start sites (TSSs) of each gene in the database. DET-specific genes had a significantly larger number of transcripts than DEGs (*p* = 6.2 × 10^-41^) (**Fig. 3C**) and also had a larger number of multiple TSSs (**Fig. 3D**) (*p* = 5.2 × 10^-13^).

**Fig. 3:**
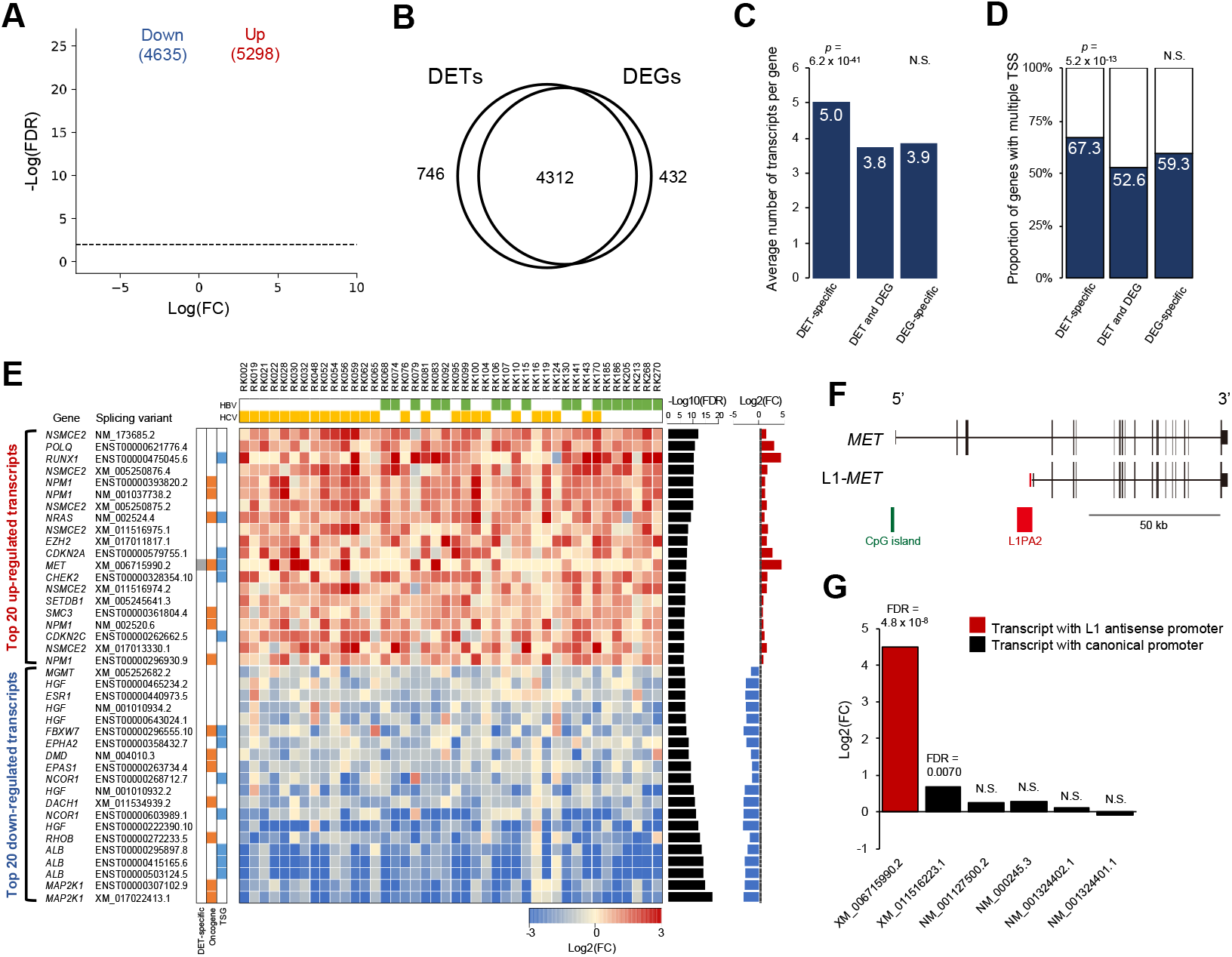
Landscape of significantly differentiated transcripts between the HCCs and the matched non-cancerous livers. (**A**) Volcano plot of the differentially expressed transcripts in the cancers and the non-cancerous livers. Red and blue dots represent transcripts that were significantly up- and down-regulated with FDR < 0.01. (**B**) Venn diagram of the number of genes with DETs and DEGs. (**C**) Comparison of the average number of transcripts per gene for each item in the Venn diagram in Fig. 3B. *P*-values for the abundance of transcripts per gene in DET-specific and DEG-specific genes were calculated by Wilcoxon rank-sum test. The numbers of the transcripts in each gene were compared for DET-specific genes vs. the others, and DEG-specific genes vs. the others. (**D**) Comparison of the proportion of genes with multiple TSS for each item in the Venn diagram in Fig. 3B. *P*-values for the enrichment of genes with multiple TSS in DET-specific and DEG-specific genes were calculated by Fisher’s exact test. The numbers of the TSS in each gene were compared for DET-specific genes vs. the others, and DEG-specific genes vs. the others. (**E**) Landscape of DETs in driver genes. Twenty up-regulated and down-regulated DETs are shown in order of lower FDR. Heatmap showing the expression differences (log2(Fold change)) in the cancer and matched non-cancerous livers. (**F**) Schematics diagram of the structural differences in L1-*MET* (XM_006715990.2) and other *MET* transcripts. L1-*MET* has the L1-derived first exon shown in red. (**G**) Bar chart showing the expression differences between the cancers and matched livers in *MET* transcripts. The red bar represents *MET* transcript with L1 antisense promoter, and black bars represent MET transcripts with the canonical promoter. *P*-values were obtained by quasi-likelihood methods by EdgeR software (Robinson et al. 2010). Benjamini-Hochberg method was used for multiple testing correction (FDR). N.S.: Not significant.

Among the 9,933 DETs, 200 transcripts (99 genes) were from previously reported driver genes (**Fig. 3E**; **Supplemental Table S9**; Bailey et al. 2018; Ding et al. 2017; Fujimoto et al. 2016), and 13 genes (16 transcripts) of them were DET-specific. Of these, transcripts of *MET* oncogene (XM_006715990.2, XM_011516223.1) were up-regulated in HCCs, and tumor suppressor genes such as *HLA-B* (ENST00000359635.9, ENST00000450871.6) and *APC* (NM_000038.5) were down-regulated (**Supplemental Table S9**). Transcripts from 16 HCC driver genes (*ALB*, *APOB*, *ASH1L*, *CDKN2A*, *CPS1*, *CTNNB1*, *DHX9*, *IDH1*, *IL6ST*, *NCOR1*, *NFE2L2*, *NRAS*, *NSMCE2*, *NUP133*, *SETDB1* and *XPO1*) were significantly differentiated (**Supplemental Table S9**; Bailey et al. 2018; Ding et al. 2017; Fujimoto et al. 2016). Of these, transcripts from *ASH1L*, *CDKN2A*, *CTNNB1*, *DHX9*, *NRAS*, *NSMCE2*, *NUP133*, *SETDB1* and *XPO1* were up-regulated in the HCCs and others were down-regulated. *ALB* is known to be highly expressed in liver, and 3 major transcripts (ENST00000503124.5, ENST00000415165.6 and ENST00000295897.8) were detected in the liver and the HCCs (**Supplemental Table S9**).

We then focused on genes that had both significantly up-regulated and down-regulated transcripts (referred to as bi-directionally significantly expressed (BiExp) genes). Our analysis identified 80 BiExp genes (270 transcripts) (**Supplemental Table S10**). Among them, *AFMID*, which was a previously reported oncogenic BiExp gene (Lin et al. 2018), was successfully detected in our analysis (**Supplemental Fig. S2A**). In the 746 DET-specific genes, 42 were BiExp genes. As expected, DEGs had a smaller proportion of BiExp genes than that of DET specific genes (*p* = 1.6 × 10^-15^) (**Supplemental Table S11**), as BiExp genes had both up- and down-regulated transcripts and their expression levels canceled out each other.

### Exonization of transposable elements (TEs) in the HCCs

A transcript of *MET* (XM_6715990.2) was ranked first in the significantly up-regulated in the HCCs among DET-specific driver genes (FDR = 4.8 × 10^-8^) (**Fig. 3E**). A previous study reported that the first exon of XM_006715990.2 was L1-derived, and this transcript is expressed by the L1 antisense promoter (referred to as L1-*MET*) (**Fig. 3F**; Hur et al. 2014). L1-*MET* was remarkably up-regulated in the HCCs compared to the other MET transcripts with the canonical promoter (**Fig. 3G**). Since exonization of TEs were observed in the HCCs and livers (**Fig. 2B)**, we next focused on cancer-specific TE-derived exons (n = 829) (**Supplemental Table S12**). We found that they were significantly enriched in the first and last exons (*p* = 7.6 × 10^-43^ for the first exon, *p* = 3.9 × 10^-7^ for the last exon) (**Supplemental Fig. S2B,C**). Analysis of repeat families showed that L1 sense and antisense, L2 antisense, ERVL sense and antisense, ERVL-MaLR sense, and LTR sense were significantly enriched in the first exon of cancer-specific transcripts (**Supplemental Table S12**). The liver-specific TE-derived exons were also significantly enriched in the first and last exons (*p* = 2.9 × 10^-6^ for the first exon, *p* = 0.0012 for the last exon) (**Supplemental Fig. S2B,C**), and by repeat families, the L2 antisense was significantly enriched in the first exon (*p* = 8.6 × 10^-5^) (**Supplemental Table S13**).

### Expression from HBV

We then analyzed reads containing the HBV genome sequence in 18 HBV-positive patients. Of these samples, 13 HCCs and 10 matched non-cancerous livers expressed transcripts having HBV genome sequence (HBV transcript) (**Supplemental Fig. S4**). We classified transcripts having the HBV genome sequence into HBV-only and HBV-human fusion transcripts. We estimated transcriptional start and end sites based on poly-A sequences in each transcript, and analyzed these locations on the HBV genome. Most of the TSSs of HBV transcripts were clustered in the upstream region of the PreS2 gene (around 1 bp – 100 bp in the HBV reference genome) (**Fig. 4A**; **Supplemental Fig. S4**) as shown in a previous study using the CAGE method (Altinel et al. 2016). We then compared the expression levels of the transcripts with HBV sequences. Expression levels of each HBV transcript were significantly different between the HCCs and the livers (*p* = 0.046) (**Fig. 4B**). However, comparison of the total amounts of HBV transcript was not significantly different between the HCCs and the matched livers (**Supplemental Fig. S5A**), suggesting that transcript-level analysis with long-reads can detect significantly differentiated transcripts from HBV.

**Fig. 4:**
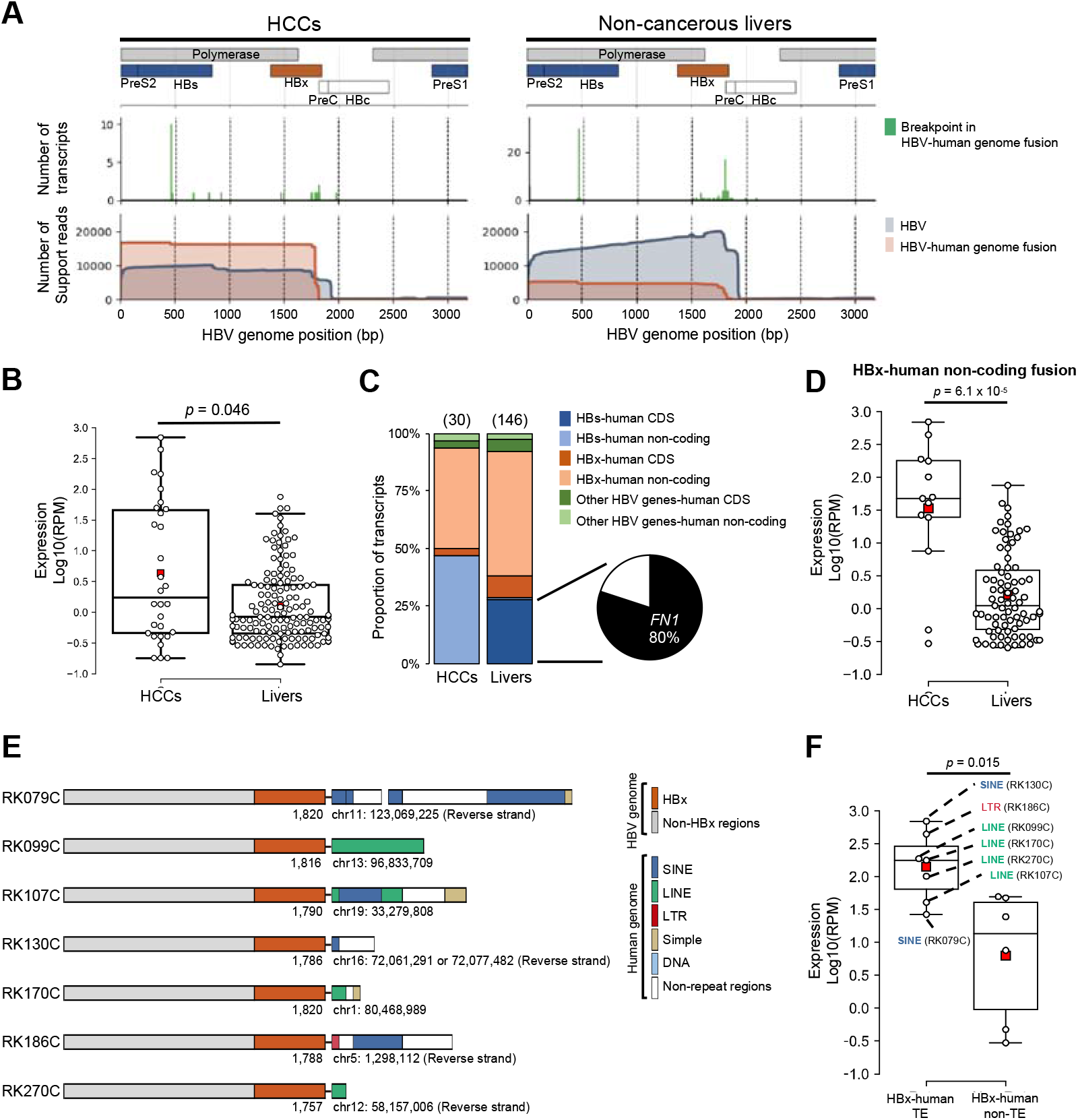
Identification of HBV-human genome fusion transcripts. (**A**) Visualization of RNA-seq coverage in the HBV genome region in the HCCs and the non-cancerous livers. (Upper panels) Schematic diagram of HBV genome structure. (Middle panels) Bar chart of breakpoints in HBV-human fusion transcripts. (Lower panels) Coverage of HBV transcripts (HBV alone) and HBV-human fusion transcripts are shown in blue and red. (**B**) Comparison of expression levels of HBV-human fusion transcripts in the HCCs and the livers. Transcript abundance was measured in reads per million reads (RPM) and log10 converted values for RPM+1 were shown in the boxplot. *P*-values were calculated by Wilcoxon rank-sum test. (**C**) Comparison of the proportion of HBV-human fusion transcripts in the HCCs and livers. The stacked bar chart shows the distribution (%) of detected transcripts by type. The total number of transcripts is shown above the bars. The pie chart shows the proportion of HBs-*FN1* fusion transcripts in HBs-human CDS in the livers. (**D**) Comparison of expression levels of HBx-human non-coding fusion transcripts in the HCCs and livers. *P*-values were calculated by Wilcoxon rank-sum test. (**E**) Schematic diagram of HBx-human TEs fusion transcripts. (**F**) Comparison of expression levels in HBx-human TE fusion and HBx human non-TE fusion transcripts in HCCs. *P*-values were calculated by Wilcoxon rank-sum test.

Thirty and 146 HBV-human fusion genome transcripts were detected in the HCCs and the livers (**Supplemental Table S14**). Breakpoints of HBV-human fusion transcripts were clustered in the HBs gene region (450 bp – 470 bp) and downstream of the HBx gene region (1750 bp – 1840 bp) (middle panel of **Fig. 4A**). Therefore, we focused on HBs and HBx genes, and classified HBV-human fusion transcripts into “HBs-human CDS fusion”, “HBs-human non-coding fusion”, “HBx-human CDS fusion”, “HBx-human non-coding fusion”, “Other HBV genes-human CDS fusion”, and “Other HBV genes-human non-coding fusion” (**Fig. 4C**). Forty HBs-human CDS fusion transcripts were detected in the livers, but not detected in the HCCs (**Fig. 4C**). Among them, 32 (80.0 %) were HBs-*FN1* fusion transcripts (**Fig. 4C**; Furuta et al. 2018). Expression levels were then compared between the HCCs and livers in these categories (**Fig. 4D**; **Supplemental Fig. S5B**). HBx-human non-coding fusion showed higher expression levels in the HCCs than that in the livers (*p* = 6.1 × 10^-5^) (**Fig. 4D**), however, other categories did not show significant differences (**Supplemental Fig. S5B**). Previous studies reported that LINE1(L1), a retrotransposon, forms HBx-L1 fusion transcript, promotes epithelial-mesenchymal transition (EMT)-like changes, and induces carcinogenesis in HCC (Lau et al. 2014; Liang et al. 2016). We thus next focused on HBx-human TE fusion and identified 4 HBx-LINE fusion transcripts, 2 HBx-SINE fusion transcripts, and 1 HBx-LTR fusion transcript (**Fig. 4E**; **Supplemental Fig. S5C,D**). The expression level of HBx-human TE fusion was significantly higher than that of HBx-human non-TE fusion in the HCCs (*p* = 0.015) (**Fig. 4F**; **Supplemental Fig. S5E**).

### Fusion genes

We detected 164 cancer-specific fusion transcripts (**Supplemental Table S15**). Although recurrent fusion event of the same gene pairs was not detected, six genes, *SENP6*, *TBRG1*, *SDC2*, *ABCD3*, *LGSN*, and *LDLR* appeared recurrently, but fused with different partners (**Fig. 5A**). We performed experimental validation with RT-PCR, and all candidates (12/12) were successfully validated (**Fig. 5B**; **Supplemental Table S16**). Within the samples of the current study, 15 have been subjected to fusion gene detection by short-reads in the ICGC pan-cancer study (ICGC/TCGA Pan-Cancer Analysis of Whole Genomes Consortium 2020). A comparison between the short- and long-reads revealed that 41 out of 50 fusions (82.0 %) were not detected in the short-reads and only 9 fusions (8.0 %) were commonly detected (**Fig. 5C**). The expression levels of the 9 commonly detected fusion genes were significantly higher than those of the 41 fusion genes detected only by the long reads (*p* = 0.00014) (**Supplemental Fig. S6A**). This comparison suggests that many unidentified fusion genes exist in the short-reads studies, and a combination of short- and long-reads can detect more fusion genes. We also compared the fusion genes and structural variations (SVs) detected by the previous whole-genome sequencing (Fujimoto et al. 2016; Fujimoto et al. 2012). Among the 164 fusion transcripts, 21 had SVs that can cause the fusion transcripts, and their expression levels were significantly higher than others (*p* = 4.2 × 10^-6^) (**Fig. 5D**; Fujimoto et al. 2016).

**Fig. 5:**
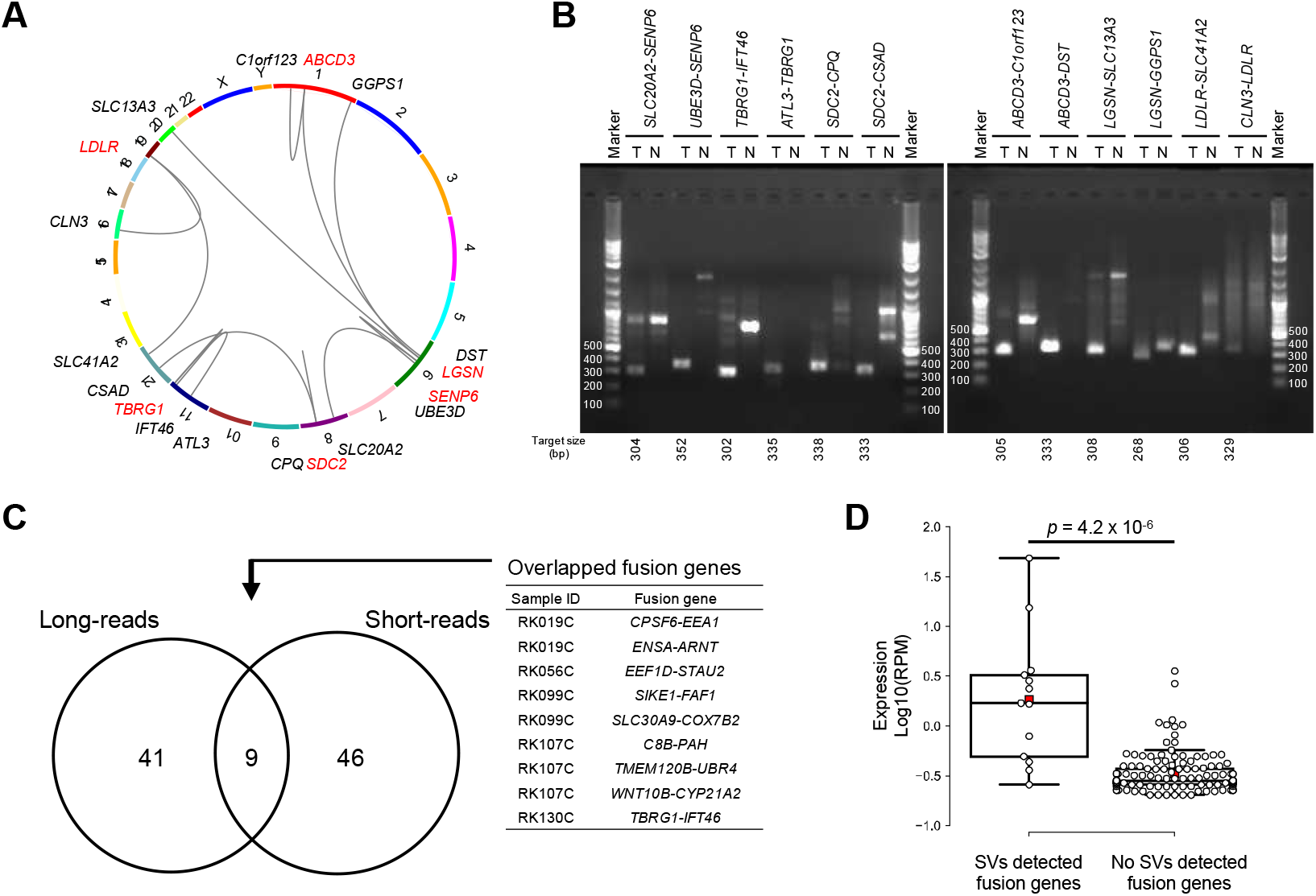
Identification of fusion genes. (**A**) Circos plot of recurrent fusion genes. Genes shown in red are recurrent fusion genes. (**B**) PCR validation of recurrent fusion genes. The results of electrophoresis of each fusion gene in the cancer (T) and matched non-cancerous livers (N) are shown. (**C**) Venn diagram of fusion genes detected by the long-reads and the short-reads sequencers. We compared 15 HCC samples in which short-reads RNA-seq was analyzed in the ICGC study (ICGC/TCGA Pan-Cancer Analysis of Whole Genomes Consortium 2020). (**D**) Comparison of expression levels between fusion genes supported by structural variations in WGS and others. *P*-values were calculated by Wilcoxon rank-sum test.

### Comparison of expression profiles of HBV- and HCV-related HCCs

Our non-cancerous liver samples have hepatitis by infection with HBV and HCV. To examine the influence of HBV and HCV on hepatitis and HCC, we performed a clustering analysis for 16 HBV-related, 24 HCV-related, and 2 both-infected samples in the HCCs and the non-cancerous livers, respectively (**Fig. 6A**). In the livers, most HBV- and HCV-related tissues were in separate clusters, whereas the HCCs were not separated (**Fig. 6A**). Next, we analyzed genes with significantly differentiated transcripts between HCV and HBV samples. In the HCV- and HBV-related livers, 284 and 68 transcripts were significantly up-regulated in the HCV- and HBV-related livers (**Fig. 6B,C**). Highly-expressed transcripts in the HCV-related livers were significantly enriched in immune system-related pathways, especially in the interferon signaling pathway (**Fig. 6D**; **Supplemental Table S17**). Transcripts up-regulated in the HBV-related livers had no significantly enriched pathways. In the HCV- and HBV-related HCCs, we performed the same analysis, and found that immune system-related pathways were slightly enriched in differentially expressed transcripts only in the HCV-related HCCs, but not as pronounced as in the livers (**Fig. 6C**; **Supplemental Table S17,18**). In comparison between the HCCs and the livers by virus, genes of cell cycle-related pathways were significantly up-regulated in the both HBV- and HCV-related HCCs as observed on all HCCs (**Fig. 6D**; **Supplemental Table S8,19,20**).

**Fig. 6:**
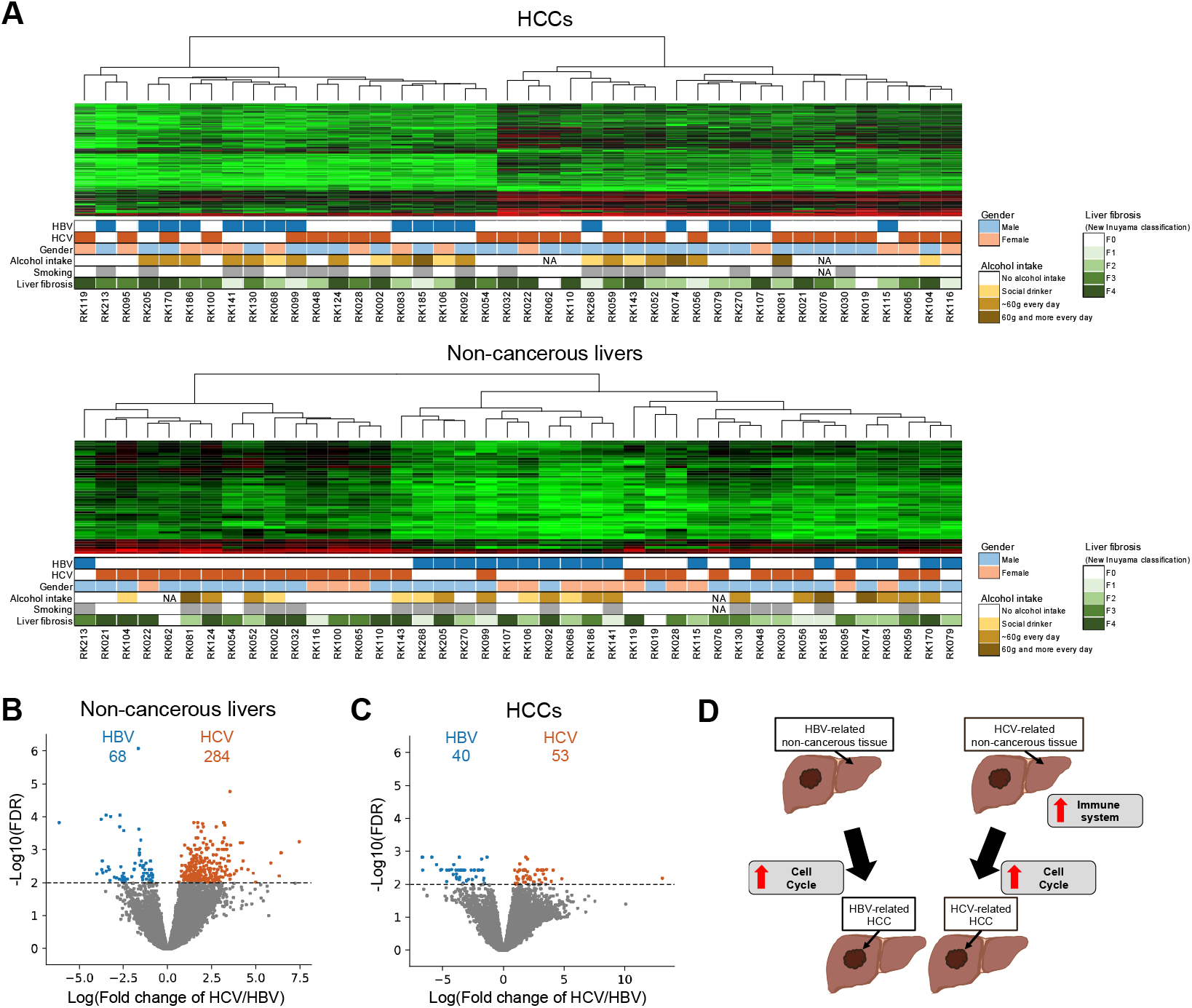
Comparison of expression profiles of HBV-related and HCV-related HCCs. (**A**) Clustering by the transcriptome of 16 HBV-related tissues, 24 HBV-related tissues and 2 both-related tissues in the HCCs (upper) and matched non-cancerous livers (lower), respectively. Clinical and pathological information of the samples are shown in the lower panels. (**B,C**) Volcano plot of the differentially expressed transcripts in HBV- and HCV-related tissues. Light blue and orange dots represent transcripts that were significantly up-regulated in HBV- and HCV-related tissues with FDR < 0.01. (**B**) Comparison in the non-cancerous liver samples. (**C**) Comparison in the HCC samples. (**D**) Schematic diagram of the expression profiles of HBV- and HCV-related tissues.

## Discussion

In this study, we used a long-read sequencer (Oxford Nanopore) for a comprehensive analysis of transcriptional abnormalities for HCCs and non-cancerous livers. We first developed an analysis pipeline (SPLICE) to analyze full-length transcripts (**Fig. 1A,B**). SPLICE removed possible errors around splicing junction sites and pseudo fusion transcripts by considering alignment and mapping errors (**Fig. 1B**). Comparison with short-reads showed high concordances in the estimation of gene expression level, suggesting that our analysis was reliable (**Supplemental Fig. S1F**). In the analysis of the HCCs and the livers, 15.7 % and 13.2 % of transcripts were not found in the database and considered novel (**Fig. 1D**). The expression levels of the novel transcript were significantly lower than those of known transcripts, and deeper sequencing would detect a larger number of novel transcripts (**Fig. 1E,F**).

Our analysis identified novel exons, and the majority of them were first and last exons (**Fig. 2A**). The novel first exons can be produced by TSSs from unknown alternative promoters and may contribute to tissue-specific or disease-associated mRNA regulation. Novel last exons can cause new UTRs and may affect the stability of mRNA. Since the average conservation scores of the novel exons were lower than those of known exons, most of them would not have important functions (**Fig. 2D**). However, some novel exons were highly conserved and would have essential functions for biological activities (**Fig. 2D,E**). In the *MYT1L* gene, a highly-conserved novel exon was detected (**Fig. 2E**), and this transcript was significantly down-regulated in the HCCs (**Supplemental Fig. S3A**). Previous studies reported that expression of *MYT1L* is neuron-specific (Mall et al. 2017), and has been suggested to play a critical role in neuronal differentiation and maintenance (Mall et al. 2017; Matsushita et al. 2014). However, our analysis identified a new transcript with the novel exon of this gene in the liver. Several studies suggested that *MYT1L* suppresses proliferation of glioblastoma and this transcript would have tumor-suppressive effects in HCCs as well (Hu et al. 2013; Melhuish et al. 2018).

This study also identified the exonization of transposable elements (TEs). Interestingly, 30-50 % of the novel exons in protein-coding genes were derived from TEs (**Fig. 2B**), suggesting that TEs can be a source of functional diversity of genes (Cordaux and Batzer 2009). Strands of the exonized TEs were not random and patterns were different among the location of the exons (**Fig. 2C**); first and middle exons had higher proportions of TEs encoded on opposite (antisense) strands, whereas the opposite pattern was observed in TEs of last exons (**Fig. 2C**). A previous study suggested that antisense of LINE1 has a promoter function and can start transcription (Hur et al. 2014; Speek 2001). This can partly explain the strand bias of TEs of first exons. The strand bias of TEs of the middle exon can be caused by the asymmetric distribution of TEs in introns. TEs on sense strands generate harmful poly-A signals in introns and this results in the reduction of TEs in sense strands in introns (Roy-Engel et al. 2005; Smit 1999). *Alu* has polyadenylation signals in the sense strand, and this can generate last exons, which can explain the strand bias of *Alu* of the last exon (**Fig. 2C**; Chen et al. 2009).

Comparison of expression level between HCCs and non-cancerous livers showed differentiation of several transcripts with TE-derived exons. A transcript with TE-derived exon from an important oncogene, *MET*, (L1-*MET*) was significantly overexpressed in HCCs (**Fig. 3F,G**). L1-*MET* was reported to be involved in the progression of colorectal cancer and bladder cancer (Hur et al. 2014; Wolff et al. 2010), suggesting that it plays an important role in HCCs as well. Since L1-*MET* is suggested to be regulated by the L1 antisense promoter (**Fig. 3F**; Hur et al. 2014; Speek 2001; Wolff et al. 2010), its expression is independent of that of other *MET* transcripts. Indeed, 6 transcripts were observed in *MET*, however, only L1-*MET* was highly overexpressed in the HCCs (**Fig. 3G**). We also found that L2-*HRH1* was significantly upregulated in the HCCs, and the expression level of the transcript was not correlated to other transcripts in the gene (**Supplemental Fig. S2D,E**). *HRH1* gene is known to promote the progression of HCC and this transcript would have an oncogenic role (Zhao et al. 2020). TE fusion transcripts (HBx-TE fusion) were also identified in HBV-related HCCs and livers (**Fig. 4F**). Previous studies have shown that the expression of HBx-LINE1 affects the transactivation of β-catenin and is involved in the activation of Wnt signaling (Lau et al. 2014; Liang et al. 2016). As overexpression of HBx results in HCCs progression (Koike et al. 1994), the other HBx-TE fusion transcripts can be involved in cancer progression with detrimental functions, although functional analysis is required to confirm their roles.

Most RNA-seq studies identified differentially expressed genes (DEGs), and a lot of differentially expressed transcripts (DETs) are likely to have been overlooked. In this study, we detected 9,933 DETs (**Fig. 3A**). Seven hundred forty-six genes with DETs were not detected by the gene-level analysis (DET-specific genes), indicating that analysis of transcripts can detect novel driver genes (**Fig. 3B**). Our analysis also showed possible mechanisms of regulation of splicing variants. DET-specific genes had a higher number of transcripts per gene and a higher proportion of transcripts with multiple TSSs (**Fig. 3C,D**). This suggests that multiple transcripts regulated by different promoters canceled out each other, and thus expression differences were not detected in the gene-level analysis. We also found 80 genes with both significantly up-regulated and down-regulated transcripts (BiExp genes) (**Supplemental Table S10**). In addition to the previously reported *AFMID* (Lin et al. 2018), genes related to aberrant cell proliferation (*CEACAM1*, *CASP8*, *LDHB*, *SPINT2*) (Abou-Rjaily et al. 2004; Boege et al. 2017; Brisson et al. 2016; Morris et al. 2005) were detected as BiExp genes (**Supplemental Table S10**). Each transcript in BiExp genes would be regulated differently and have different functions.

We then compared the expression level of transcripts in the HCCs and the non-cancerous livers between HBV and HCV (**Fig. 4**). Clustering analysis did not clearly separate HBV and HCV HCCs, whereas the livers were clustered into three groups; HCV cluster, HBV cluster and cluster with both types (**Fig. 4A**). This suggests that the difference in virology mainly affects the phenotype of the livers (**Fig. 4D**). Pathway analysis for genes differentiated between HCV- and HBV-livers showed enrichment of immune-related genes (**Supplemental Table S17**). This result is consistent with previous studies (Iizuka et al. 2002; Sun et al. 2019) and suggests that HBV and HCV may induce different expression changes, but similar mechanisms work after cancer development (**Fig. 6A**; **Supplemental Table S17,18,19,20**).

Although we obtained many interesting discoveries by analyzing full-length transcripts, our study has limitations that should be addressed in future studies. First, our analysis did not determine the structure of the entire CDS for 38.3 % of mapped reads. The majority of the incomplete transcripts lacked 5’ regions (**Supplemental Fig. S1C,D**), and this can be explained by the process of cDNA synthesis and degradation of RNA samples. cDNA synthesis starts from 3’ends and therefore, the inactivation of reverse transcriptase or degradation of RNA samples result in cDNA lacking 5’ ends (**Supplemental Fig. S1E**). This mechanism can also affect fusion-gene detection (**Fig. 5**; **Supplemental Fig. S6B,C**). Improvement of cDNA synthesis and the use of high-quality RNA can increase the rate of full-length transcripts. Second, our analysis removed the change of splicing sites within 5 bp to remove alignment errors (**Fig. 1B**). We consider that this cutoff value is necessary due to currently available high-error reads (**Supplemental Table S4**). However, sequencing technologies and basecallers are improving, and in the near future, we should be able to use a smaller cutoff value and identify larger numbers of splicing changes. In this study, we developed the SPLICE pipeline for transcriptome analysis for a long-read sequencer. Validation with MCF-7 cell line and PCR showed the reliability of our analysis. Using this method, we analyzed full-length transcript of HCCs. We detected 746 genes with DETs, which should be difficult to detect by short-read sequencers. We also detected cancer-related genes with TE-derived exons. In the analysis of fusion genes, we found 6 recurrent genes that fused with other genes. Additionally, by comparing HBV-with HCV-related livers, significant differentiations in immune-related genes were detected. To our knowledge, this is the first report about transcript aberrations in cancer clinical samples using a long-read sequencing technology. Our result strongly suggests that direct observation of transcripts with long-reads contributes to understanding the true picture of transcript aberration in cancer.

## Methods

### Samples

RNA from HCCs and adjacent non-cancerous livers from 42 patients was used for this study. These samples have been used for short-read sequencing and reported by several previous studies (**Supplemental Table S1**; Fujimoto et al. 2016; Fujimoto et al. 2012; ICGC/TCGA Pan-Cancer Analysis of Whole Genomes Consortium 2020), and list of somatic mutations (point mutations, short indels and structural variations), gene expression level and fusion gene by short-read analysis are available (Fujimoto et al. 2016; ICGC/TCGA Pan-Cancer Analysis of Whole Genomes Consortium 2020). RNA sample qualities were evaluated with RIN (RNA Integrity Number) values by Bioanalyzer (Agilent), and samples with high RIN values ≥ 8 were selected. RNA from 16 HBV-related HCCs, 24 HCV-related HCCs, 2 HBV and HCV-related HCCs and their matched non-cancerous livers were sequenced.

### Library preparation and cDNA sequencing

Full-length cDNA was synthesized from total RNA (1 μg) by SMARTer^®^□ PCR cDNA Synthesis kit (Clontech) according to the manufacture’s instruction. For primer digestion, cDNA samples were treated by exonuclease I (NEB) at 37 °C for 30 min. After purification with Agencourt AMPure XP magnetic beads (Beckman Coulter), libraries were constructed using the Ligation Sequencing Kit (SQK-LSK109) (Oxford Nanopore) and sequenced with SpotON FlowCell MK I (R9.4) (Oxford Nanopore) according to the manufacture’s protocol. Basecalling from raw data (FAST5 format) was performed using the Guppy software (version 3.0.3) (Oxford Nanopore).

### Analysis pipeline

We constructed an analysis pipeline named SPLICE (**Fig. 1A**). Low-quality reads (average base quality < 15) were filtered out. Reads were mapped to the reference genome sequence (hg38) and the reference transcriptome sequence (GENCODE (version 28) and RefSeq (release 88)) with minimap2 software (Li 2018). For reads mapped to the reference genome sequence, ≧ 60 bp unmapped region (soft-clipping regions) was re-mapped to the reference genome sequence (hg38). Since sequencing errors near splicing junctions can lead to errors in predictions of splicing sites, we compared the mapping results to the reference genome and the reference transcriptome sequence, and removed reads if the number of matches to the reference genome sequence was smaller than that to the reference transcriptome sequence.

### Identification of known transcripts

Genomic locations of known transcripts were obtained from GENCODE (version 28) and RefSeq (release 88), and transcripts that were commonly registered in both annotations were unified to GENCODE transcript IDs (**Fig. 1A**). We assigned each exon in the transcripts to the exons in the database if the positions of the splicing junction sites at both ends of the transcript’s exon and reference exon were perfectly matched. We then considered the combination of the exons and transcript ID in the database were assigned. Transcripts that were not found in the database were classified as novel. For the annotation, we did not consider UTRs length for the classification, because qualities of reads at the ends of reads were low. Reads that contained all coding exons of transcripts in GENCODE or RefSeq were classified to known “full-length CDS”. When reads had a part of exons of a known transcript but do not cover all exons, they were classified to known “partial-length CDS”. Among partial-length CDSs, if the reference transcripts had 2 or more exons and reads had only one exon, we removed the candidates as highly truncated reads. For partial-length CDSs reads annotated to multiple known transcripts, we annotated them to the longest known transcript among matched known transcripts. After the annotation, we removed transcripts if their proportion of expression level of a transcript was less than 1 % of the total amount of expression of the gene.

### Permutation test

We performed a permutation test to examine the skew of the distribution of repeat types and strands. Random sequences corresponding to novel exon lengths were extracted from within the gene regions, and overlapping repeat types and their strands were obtained. This procedure was repeated 1,000,000 times, and the distributions of repeat types and their strands were used as null distribution. Adjustment of the multiple testing was performed by Bonferroni’s correction.

### Identification of novel transcripts

Reads that were not annotated as known transcripts were classified as “novel exon length”, “novel exon combination” or “novel exon” (**Fig. 1C**). “Novel exon length” reads contain known exons with different lengths from the reference database. As described above, change of splicing junction sites ≥ 5 bp was considered as length change. If splicing junction sites changed < 20 bp for a known splicing junction site, we evaluated the mismatch rate within ± 5 bp regions from splicing junction sites and removed candidates if error rates were ≧ 20 %. Novel exons were defined as expressed regions that were not overlapped with known exons. Novel exon candidates were removed if they were expressed only in a single novel exon. We removed candidates if their proportion of transcript expression level was < 1 % of the total amount of expression level of the gene.

### Identification of fusion genes

Reads mapped to two protein-coding genes were considered as fusion gene candidates. Possible errors in fusion gene candidates were filtered out as follow: (1) Mapping errors due to highly homologous genomic regions can cause fusion gene candidates, therefore, we compared the results of mapping to the reference genome and the reference transcriptome sequences and removed candidates if both were inconsistent (**Fig. 1B**). (2) Fusion gene candidates can be generated by mis-ligation of two transcripts during library preparation, we therefore removed candidates that contained primer sequences for cDNA synthesis between two genes (**Fig. 1B**). (3) Artificial chimeric reads may be generated from highly expressed genes, because of the higher chance of mis-ligation in library preparation, thus, we removed candidates if their proportion of transcript expression level was < 1 % of the average expression level of both genes. (4) Candidates with low expression levels (support reads < 3) in 2 or more samples in the matched non-cancerous livers were removed as artifacts. (5) Candidates were removed as read-through transcripts if the two transcripts were the same strand and were in close proximity to each other in the genome (< 200,000 bp).

We further compared the fusion genes with structural variations (SVs) detected by the previous whole-genome sequencing (Fujimoto et al. 2016; Fujimoto et al. 2012). If the breakpoints of an SV were located in intron or exon regions of the fusion gene, it was determined to be an SV that could cause the fusion gene.

### Comparison of gene expressions between the HCCs and the non-cancerous livers

To compare expression levels in the HCCs and the matched non-cancerous livers, we used edgeR software (Robinson et al. 2010). The number of mapped reads for each transcript was normalized using trimmed mean of M value (TMM) normalization method (Robinson and Oshlack 2010). Differential expression analysis was performed using the quasi-likelihood method (Wedderburn 1974). The Benjamini-Hochberg method was used for the multiple testing correction (Benjamini and Hochberg 1995).

### Clustering analysis

To compare the expression profiles in HBV- and HCV-related tissues, we applied k-means clustering to the RNA-seq data. For the log10 converted value of TMM normalized RPM+1, we filtered out candidates with a variance < 0.1. We performed k-means clustering, and then excluded candidates that were not involved in the clustering by the variance analysis (p-value < 1 × 10^-5^).

## Data access

The source code of SPLICE is freely available from Github (https://github.com/hkiyose/SPLICE). Gene expression data of MCF-7 cells by short-read sequencer was obtained from ENCODE website (https://www.encodeproject.org/files/ENCFF629VUS/). <u>Conservation score of 100 vertebrates was downloaded from UCSC genome browser</u> (http://hgdownload.cse.ucsc.edu/goldenPath/hg38/phyloP100way/). The sequencing data have been deposited in the National Bioscience Database Center (NBDC) under accession XXXX.

## Acknowledgements

This work was supported by Grant-in-Aid for Scientific Research (B) from Japanese Society for the Promotion of Science (18H02680 to A.F.), the Platform Program for Promotion of Genome Medicine (Grant Number 19km0405207h0004, to A.F.) and Takeda Science Foundation (to A.F.).

The super-computing resource was provided by Human Genome Center, Institute of Medical Science, the University of Tokyo. We acknowledge the staff at Laboratory for Cancer Genomics, RIKEN, for their technical assistances.

## Author Contributions

AF designed the study. HK performed the computational analyses. HK performed the experiments. HK, HN, MS and AF interpreted the results. HK, JHW and AF wrote the manuscript. All authors approved the final manuscript.

